# Structural analysis of *de novo* STXBP1 mutation in complex with syntaxin 1A reveals a major alteration in the interaction interface in a child with developmental delay and spasticity

**DOI:** 10.1101/642702

**Authors:** Ehud Banne, Tzipora Falik-Zaccai, Esther Brielle, Limor Kalfon, Hagay Ladany, Danielle Klinger, Dina Schneidman-Duhovny, Michal Linial

## Abstract

STXBP1, also known as Munc-18, is a master regulator of neurotransmitter release and synaptic function in the human brain through its direct interaction with syntaxin 1A. STXBP1 related disorders are well characterized and cover a diverse range of neurological and neurodevelopmental conditions. Through exome sequencing of a child with developmental delay, hypotonia and spasticity, we found a novel *de novo* insertion mutation of three nucleotides in the STXBP1 coding region, resulting in an additional arginine after position 39 (R39dup). Inconclusive results from state-of-the-art variant prediction tools mandated a structure-based approach using molecular dynamics (MD) simulations of the STXBP1-syntaxin 1A complex. Comparison of the interaction interfaces of the wild type and the R39dup complexes revealed a reduced interaction surface area in the mutant, leading to destabilization of the interaction. We applied the same MD methodology to 7 additional previously reported STXBP1 mutations. We find that the stability of the STXBP1-syntaxin 1A interface correlates with the reported clinical phenotypes. We illustrate a direct link between a patient’s genetic variations and the observed clinical phenotype through protein structure, dynamics, and function.

## MAIN TEXT

Normal brain function requires a highly coordinated regulation of neurotransmitter (NT) release from synaptic vesicles (SVs). The critical stage in this process is the fusion of SV with the plasma membrane in response to depolarization and calcium entry^1^. This step in NT release is orchestrated by a dynamic protein-protein interaction network. Proteins from the VAMP, SNAP-25, and syntaxin families, collectively known as SNAREs, function at a late step in membrane fusion^2^. The SNAREs initiate fusion by forming a tight SNARE complex that brings into close proximity the SV (via VAMP) and the plasma membrane (via SNAP-25, and syntaxin)^1^. Upon activation of the SNARE complex, it undergoes spontaneous zippering to create a tighter and more stable 4-helix bundle (called SNAREpin) that provides the needed energy for disrupting the SV lipid bilayer. This non-reversible event creates a fusion pore, enabling NT release and eventually neuronal communication^3^. In addition to the fundamental role of SNARE proteins in driving SV fusion, proteins that directly bind SNARE proteins (e.g., Munc-18, Munc-13, synaptotagmin) regulate numerous aspects of the SV life cycle, including, docking, priming, and fusion^4–6^.

The indispensable role of SNAREs in NT release accords with the rarity of mutations in any of these proteins^7^. However, mutations in the direct regulators of SNARE proteins exhibit a range of clinical presentations, including autism spectrum disorder (ASD), schizophrenia, and an array of other neurological disorders^8^. Although the genetic variation underlying the clinical phenotype can be directly identified, there is no straightforward approach linking the observed mutation effect with the protein stability, interactions, and dynamics of the cellular processes involved in synaptic function.

This study examines a *de novo* mutation in STXBP1 (NM_003165.3), a high-affinity binding protein of syntaxin 1 which is also known as Munc18-1 or nSec-1^9–11^. The mutation was identified in a five years old girl of Jewish Moroccan and Christian Russian origin (Figures 1A and 1B), with severe global developmental delay and hypertonicity with spasticity. Since early age, developmental delay and episodes of tremor with excessive startle were observed. Bouts of tremor with eye rolling were noticed at the age of two and a half years. Currently, at five years old, she is incapable of independent function, and needs help in all everyday tasks.

**Figure 1.**
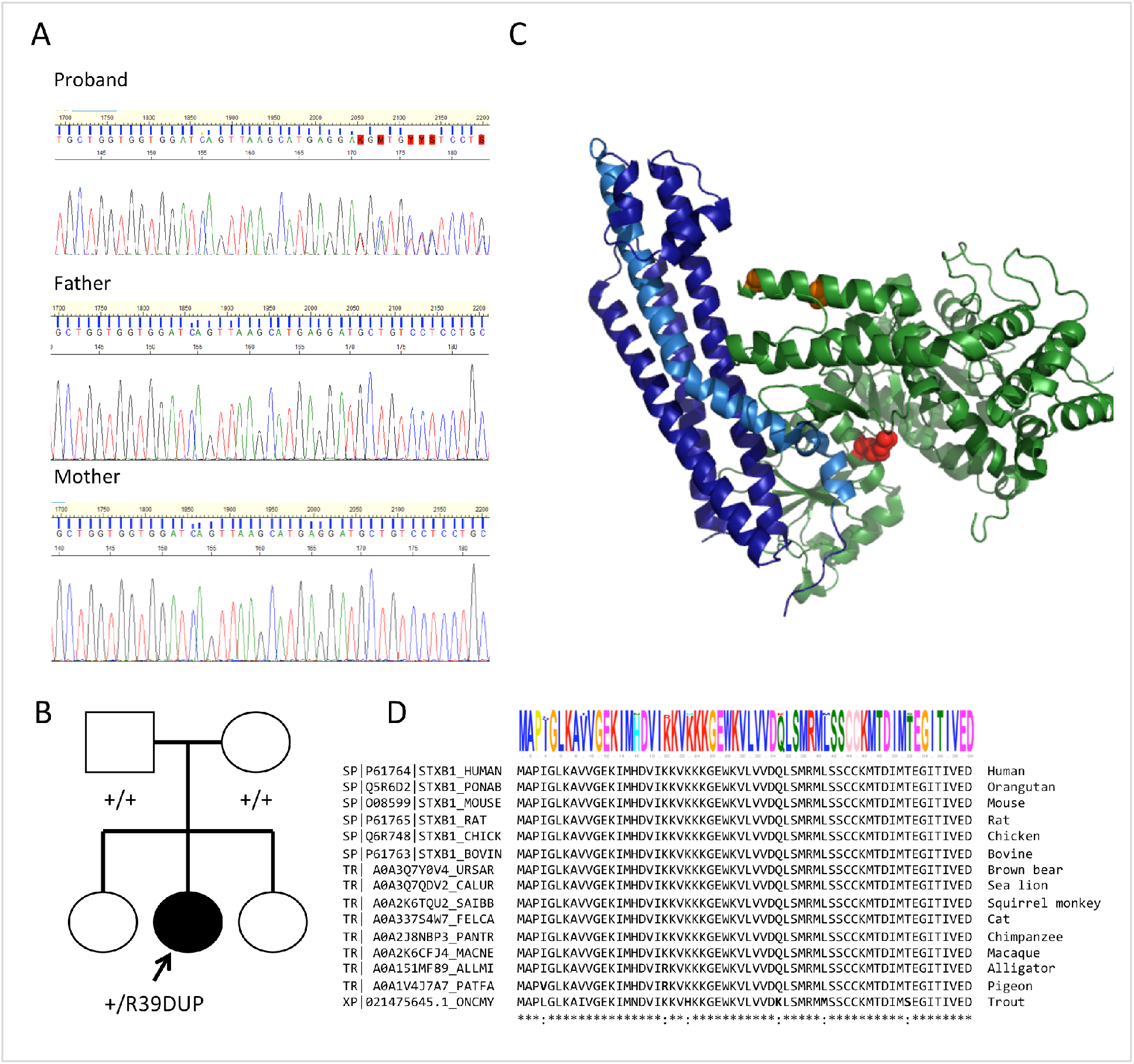
STXBP1 intragenic *de novo* variant identified in this study. (A) Sanger sequencing for the proband and the parents showing an insertion of GAG at position 116 of NM_001032221.3 transcript. (B) A family tree presentation of the DNA sequencing results shown in (A). (C) Mapping of the de novo insertion on the STXBP1-syntaxin1A protein structures. STXBP1 is colored green. The R39dup position is shown in red. The two phosphorylation sites of STXBP1 (amino acids 306 and 313) are colored orange. Syntaxin 1A is colored by a dark blue, while the H3 helix that is engaged in the formation of the SNARE complex is colored by a light blue. (D) Multiple sequence alignment covers 1-60 amino acids of human STXBP1 and 14 additional vertebrates including fish, reptile, aves and various mammals is shown. The amino acids logo diagram produced (top) demonstrates the high degree of conservation among the diverse taxa. A position that lacks a full identity among the 15 representative organisms is indicated by a column. Positions of identical amino acids are indicated by asterisks (bottom).

Her physical examination reveals hypotonia with spasticity, very slow reactions, and no dysmorphism. Her family history is negative for any neurological conditions. Brain MRI, MRS (Magnetic Resonance Imaging spectroscopy), and Electroencephalogram (EEG) were completely normal. Her genetic workup included a chromosomal microarray (Affymetrix 750k) demonstrating a normal female karyotype - arr(1-22,X)x2. A whole trio exome (parents and proband) sequencing was performed. The identified mutation is an in frame insertion in STXBP1 (transcript NM_001032221.3:c.116_118dup, NP_001027392.1:p.Arg39dup, Figures 1A and 1B). This insertion was not previously described in patients with a similar phenotype in ClinVar^12^, and is also absent from healthy individuals represented in GnomAD^13^. While there are over 300 reports of mutations along the STXBP1 coding sequence (limited to variant length of <50 nt), only one case of an in frame insertion (p.Leu257_Pro258insLeu), and four cases with in frame deletions were reported (Table 1). Interestingly, 3 of the 4 in frame deletions are implicated at the N’-terminal domain of STXBP1 in the vicinity of the reported R39dup and are associated with severe clinical conditions (Table 1).

**Table 1.**
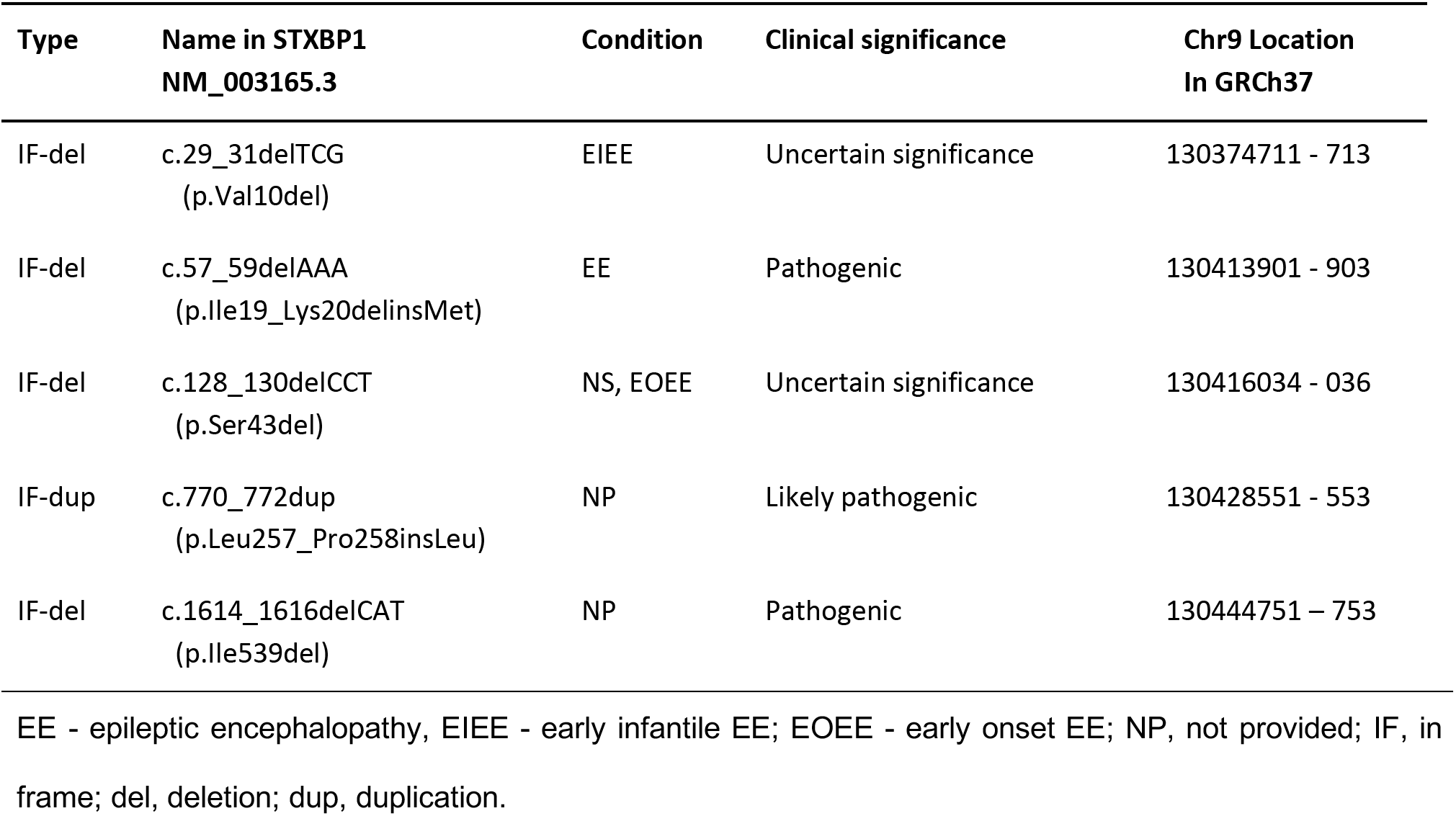
In frame indel mutations in STXBP1, according to ClinVar database (Feb. 2019)

| Type | Name in STXBP1<br>NM_003165.3 | Condition | Clinical significance | Chr9 Location<br>In GRCh37 |
| --- | --- | --- | --- | --- |
| IF-del | c.29_31delTCG<br>(p.Val10del) | EIEE | Uncertain significance | 130374711 - 713 |
| IF-del | c.57_59delAAA<br>(p.Ile19_Lys20delinsMet) | EE | Pathogenic | 130413901 - 903 |
| IF-del | c.128_130delCCT<br>(p.Ser43del) | NS, EOEE | Uncertain significance | 130416034 - 036 |
| IF-dup | c.770_772dup<br>(p.Leu257_Pro258insLeu) | NP | Likely pathogenic | 130428551 - 553 |
| IF-del | c.1614_1616delCAT<br>(p.Ile539del) | NP | Pathogenic | 130444751 - 753 |

EE - epileptic encephalopathy, EIEE - early infantile EE; EOEE - early onset EE; NP, not provided; IF, in frame; del, deletion; dup, duplication.

STXBP1 related disorders are associated with numerous neurological defects^14^. In accordance with the fundamental role of STXBP1 in NT release, the pathogenic variants in STXBP1 are now well known to cause a neurological interference of disastrous magnitude in children^15, 16^. In particular, the most severe phenotypes are early infantile epileptic encephalopathy (EIEE), Ohtahara syndrome^16^, West syndrome^17^, Dravet Syndrome^18^, and Rett syndrome^19^. In addition, STXBP1 mutations can cause milder phenotypes, including isolated intellectual disability^20^, ataxia, tremor and mental retardation without epilepsy^21^, and various manifestations of autism spectrum disorder^22^. STXBP1 pathogenic de novo truncating mutation that caused a non-syndromic intellectual disability (NSID) without epilepsy has been reported^23^. We suggest that the STXBP1 variant of R39dup underlies her medical condition.

The impact of the 3-nucleotide duplication (R39dup) on the protein function was tested according to nine state-of-the-art variant effect predictors (Table S1). Most prediction tools (e.g., VarSome^24^, PROVEAN^25^) can infer missense and frameshift mutations, but not in frame indels. Other variant effect prediction tools that allow indel assessment^26^ mark R39dup as deleterious. We anticipate that such an assignment is attributed to the extreme amino acid sequence conservation of STXBP1 across all vertebrates (Figure 1D). Consequently, it was not possible to reliably infer the clinical significance of the arginine insertion mutation. The available prediction tools apply methods for classifying variants within a single protein, but do not consider protein-protein interactions (PPI). In the case of the complex of STXBP1 and syntaxin 1A, NT release is critically dependent on the structural details of the interaction. Structure-based predictors that consider PPIs were not developed and validated for in frame indels^27, 28^. In this study, we demonstrate the pathogenicity of the R39dup variant based on the structure of the STXBP1-syntaxin 1A complex^29, 30^. We apply molecular dynamics (MD) simulations to interpret the outcome of this rare mutation in STXBP1 and provide a structure-based rationale to the observed phenotypes and the clinical outcome. We benefit from the high-resolution structures of STXBP1 and syntaxin 1A^31^ in order to investigate which contact sites predominantly present in the WT and others that are altered for the interface with the altered protein (STXBP1-R39dup).

Exonic and adjacent intronic regions were enriched from genomic DNA derived from peripheral blood via the SureSelect All Exon V5 target enrichment kits from Agilent Technologies, and paired-end sequencing was performed on illumina HiSeq4000 platforms. Raw data was uploaded to Emedgene Technologies (platform called Wells) and were aligned to the human reference genome (hg19) with BWA MEM mapping algorithm^32^. After mapping and realignment, nucleotide variants have been identified with multiple variant callers including SAMtools^33^, FreeBayes^34^, and GATK4^35^, and annotated using VEP^36^ and additional local annotations. Wells platform was used to filter out common variants and to identify candidates for a pathogenic mutation in the family after analysis and examination of the results by different inheritance modes.

The mutated structural model was generated by comparative modeling using Modeller 9.18^37^. In this template-based model, an insertion of an additional residue (R39dup) will result in helix distortion. Therefore, we computationally removed the helix (residues 35-43) from the template structure of STXBP1-interacting with syntaxin 1A (PDB: 4JEH) and replaced it with an ideal helix that includes the duplicated arginine (Figures 1A and 1B). 100 models were generated to produce the most probable protein structure, and the best scoring one according to the SOAP statistical potential^38^ was selected as a starting structure for the MD simulations.

The MD simulations were performed with GROMACS 2018 software^39^ using the OPLS-AA force field and an extended simple point charge (SPC/E) water model. Each of the mutant protein models was solvated with water molecules and ions to equalize the total system charge. The steepest descent algorithm was used for initial energy minimization until the system converged at Fmax < 1000 kJ/(mol · nm). Then water and ions were allowed to equilibrate around the protein in a two-step equilibration process. The first part of equilibration was at a constant number of particles, volume, and temperature (NVT). The second part of equilibration was at a constant number of particles, pressure, and temperature (NPT). For both MD equilibration parts, positional restraints of k = 1000 kJ/(mol · nm^2^) were applied to heavy atoms of the protein, and the system was allowed to equilibrate at a reference temperature of 300 K, or reference pressure of 1 bar for 100 ps at a time step of 2 fs. Following equilibration, the production simulation duration was 100 nanoseconds with 2 fs time intervals for the wild type (WT) or any of the mutated protein. Altogether 10,000 frames were saved for the analysis. Interaction energies between STXBP1 and syntaxin 1A were calculated for each frame of the trajectory using FoldX Suite 4.0^40^. Interaction energies are measured in kcal/mol, where more negative values correspond to a stronger interaction. In the interface contact analysis, a residue-residue contact was defined based on the inter-atomic distance, with a cutoff of 4Å.

STXBP1, through its high affinity to syntaxin 1A, plays a role in regulating NT release^41–43^. According to the available 3D structure (PDB: 4JEH), syntaxin 1A is bound in its autoinhibited, “closed” conformation state. In such a state, the SNARE domain of syntaxin is clamped in the STXBP1 central cavity. The regulatory helices of syntaxin (referred to as Habc, Figure 1C) hinder the syntaxin 1A SNARE domain and precludes its assembly into a SNARE complex.

The R39dup mutation is positioned at the interface with syntaxin 1A. We therefore compared the interaction energies of the wild type (WT) and the mutant (R39dup) in complex with syntaxin 1A. We show that the interface interaction energy for the mutant was significantly higher than that of the WT (Figure 2A). The calculated ΔG for the mutant was −16.97 kcal/mol and −23.20 kcal/mol for the WT, resulting in an average difference of 6.23 kcal/mol between the mutant and the WT protein complexes. We observed a significant structural rearrangement in the R39dup mutant interface (Figures 2B and 2C).

**Figure 2:**
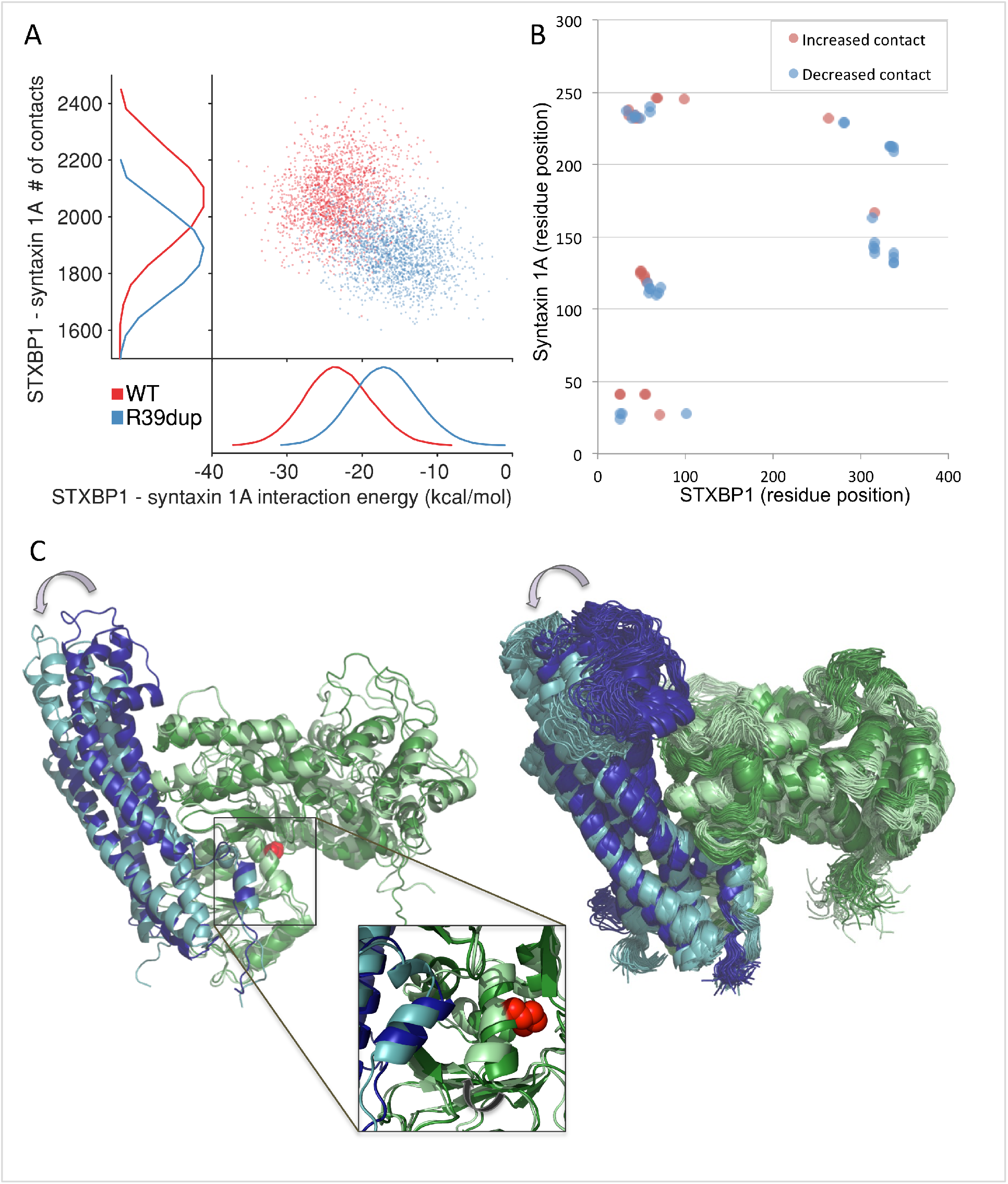
Changes in the STXBP1-syntaxin 1A interaction (WT vs. R39dup) according to MD simulations. (A) Each dot is one of the 10,0000 simulation frames of the WT (red) and the R39dup (blue). Fewer contact points are observed for the mutant (y-axis) resulting in higher interaction energy (x-axis) of the mutated R39dup protein. (B) Residue-residue contacts within 4Å that resulted in a significant alteration. An increase or decrease in the residue-residue contacts of the mutant vs. WT is shown in red and blue, respectively. (C) Structural overlay of representative frames of the STXBP1-syntaxin 1A complex. The WT is colored in dark green (STXBP1) and dark blue (syntaxin 1A), while the complex of R39dup and syntaxin are colored by light green and light blue, respectively. The arrow direction indicates the structural rearrangement of the syntaxin 1A with respect to STXBP1. A single representative frame (left) and an assemble of 50 random frames (right) are shown. In the R39dup, the helix that includes an additional arginine (red) is shifted towards syntaxin 1A (inset).

We examine what might contribute to the strong increase in the mutant interface energy. We observed that several new residue-residue contacts appear, while others disappear (e.g. residue positions 25-71, Figures 2B and 2C, Video S1). This change is attributed to the insertion of an arginine residue in the helix that results in shifting the helix residues and a re-packing of the interface with syntaxin 1A chain. In addition, we observed that the contacts in residues 280-338 mostly disappeared in the mutant with respect to the WT, suggesting that the shift in the local structural was propagating and caused a distortion further away from the position of the residue insertion. Importantly, this part of the STXBP1 protein is involved in PKC-mediated phosphorylation^44^ (Figure 1C).

We tested whether the detailed structural information of the STXBP1-syntaxin 1A complex can be used to predict additional mutations in STXBP1 according to their calculated effect on the binding interface. The clinical assignments that are reported from nine major mutation effect prediction tools are inconclusive. We tested the collection of 99 STXBP1 missense mutations reported in ClinVar and found that 42% are marked as uncertain, with an additional 8% having conflicting assignments (Figure 3).

**Figure 3.**
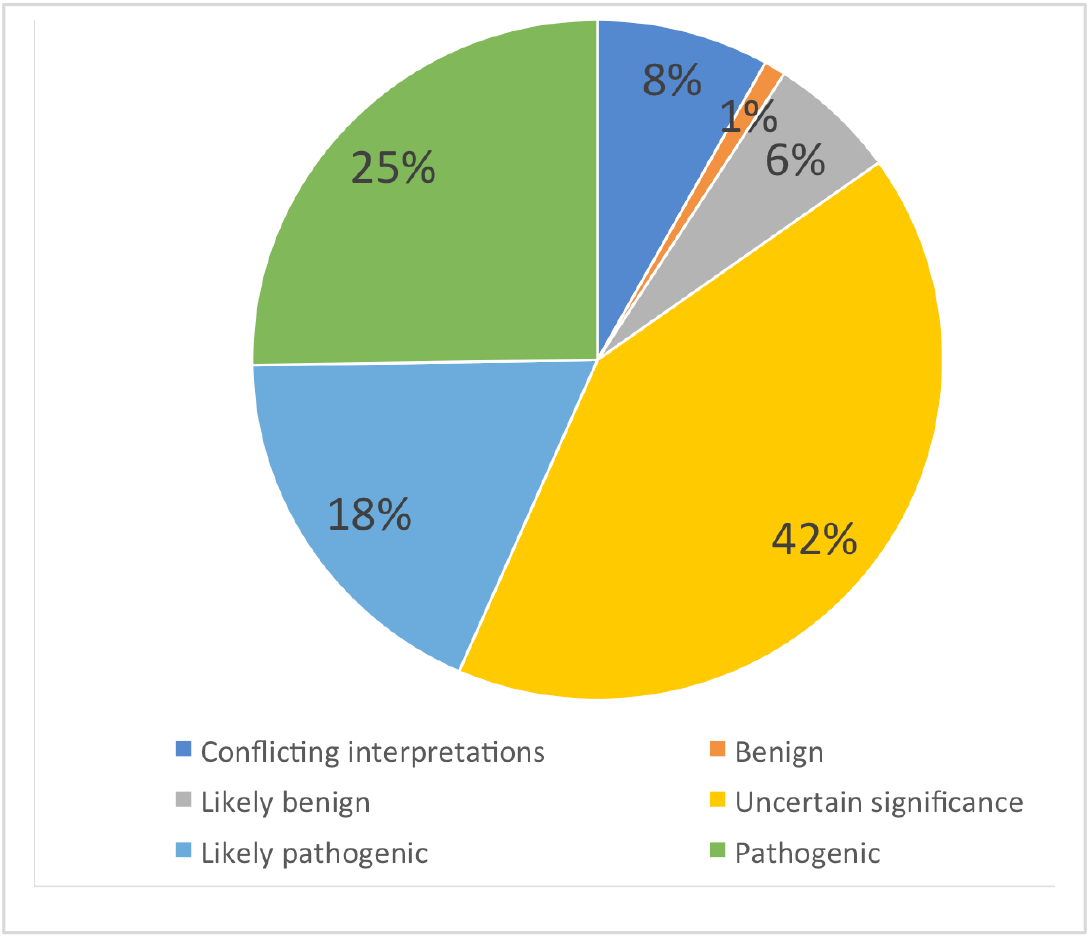
Analysis of reported 99 missense mutations in STXBP1. All 99 reported mutations from ClinVar (Feb 2019) were collected and partitioned by 6 categories for their pathological impact. Reports with conflicting results are labelled as such, and cases with no clear interpretation are marked as ‘uncertain significance’.

We then tested the applicability of the clinical assessment of the mutations’ effect using MD simulations. To this end, we applied the described MD method to seven of the missense mutations reported in ClinVar (Table 2). Six of these mutations were selected due to their proximity to the syntaxin 1A interface. An additional mutation, V84D, is remote from the interaction interface and was selected as a control. Notably, Val84 missense (V84I) is recurrent in the healthy population with an allele frequency of 5.76e-4 (GnomAD, Table 2). We find that there is an increase in the interaction energy of at least 2 kcal/mol for all tested mutations except V84D and P335L. This is explained by the fact that V84D is far from the interface with syntaxin 1A. Thus, the pathogenic outcome is likely to be attributed to other STXBP1 properties. The missense mutation P335L is labeled as uncertain significance (Table 2). Based on our results, no energetic alteration is recorded. This is in accordance with the results of single-molecule force spectroscopy showing no difference between the WT and P335L with respect to the closed state of syntaxin 1A^45^. Note that a missense mutation at the same position (P335S) is labeled as likely pathogenic, and its interaction energy is strongly affected (ΔΔG = 2.70 kcal/mol). This position is assumed to be in contact with the SNARE domain of VAMP and serves as a platform for the SNARE assembly^46^. Interestingly, an *in vitro* study showed that P335 is a critical position, and the P335A mutation still supported effective priming of SV fusion and NT release^47^.

**Table 2.** Mutations tested for interaction energy deviation from WT STXBP1.

| Mutation | ClinVar annotation, Comments <sup>a</sup> | $\Delta\Delta G$<br>(kcal/mol) | # of report in<br>GnomAD |
| --- | --- | --- | --- |
| R39dup | This report | 6.23 | - |
| R39C | In vitro, overexpression in PC12<sup>48</sup> | 1.88 | - |
| S42P | Pathogenic | 3.20 | - |
| V84D | Pathogenic, EIEE | 1.30 | 163 of 282,852 |
| P139L | Pathogenic / likely pathogenic, EIEE | 3.64 | - |
| K333E | Uncertain significance | 2.95 | - |
| P335S | Likely pathogenic | 2.70 | - |
| P335L | Uncertain significance, force spectroscopy<sup>45</sup> | -0.30 | - |

<sup>a</sup> EIEE, early infantile epileptic encephalopathy 4

In this study we present a structure-based approach of MD analysis for measuring the impact of missense and in frame mutations in the coding region of STXBP1. We applied the MD method to infer the functional outcome of a rare de novo mutation of a 3-nucleotide insertion in the coding region of STXBP1. This novel mutation resulted in the addition of an arginine between residues 39 and 40 of STXBP1, and is associated with a global developmental delay and hypertonicity with spasticity. The severity of the clinical manifestation in patients with a mutated STXBP1 gene (Figure 1) suggests that STXBP1 plays a fundamental role in neuronal communication. A comprehensive analysis of the genetic spectrum of STXBP1 (147 patients^14^) failed to detect the statistical significance of the phenotype–genotype relationships regarding all clinical aspects, including age of seizure onset, level of intellectual disability (ID), type of mutation, and more. Together with the inconsistency of in silico prediction tools (Table S1, Figure 3), it is expected that a phenotypic explanation for STXBP1 mutations should include a detailed 3D structural analysis, including the mutation’s effect on the protein-protein interactions in a presynaptic context.

The in frame R39dup is positioned at the interface of STXBP1 with syntaxin 1A, specifically at the vicinity of the core SNARE domain that is engaged in SNAREpin formation. This strategic position of the mutated protein suggests that the pathogenic outcome is a result of the alteration in the interaction of the two proteins (Figures 2A to 2C). Additionally, overexpression of mutated STXBP1 in which R39 was replaced by cysteine (STXBP1-R39C) resulted in a change in the exocytotic release probability in adrenal chromaffin and PC12 cells. In both types of cells, the binding affinity to syntaxin 1A of the mutated protein was reduced by about five-fold in comparison to the WT STXBP1^48, 49^. These *in vitro* results are consistent with our calculated changes in the interface energetics of the STXBP1-syntaxin 1A complex (Table 2). We propose that a comparative analysis based on MD simulations for calculating the deviation in protein-protein interaction energy, be applied to other complexes for which high resolution X-ray structures are available.

The broad spectrum of clinical outcomes that are associated with mutated STXBP1 argues for a global brain development defect. By manipulating the expression of STXBP1 in the developing mouse brain and introducing pathological versions of the gene, it was shown that STXBP1 dominates the radial migration of cortical neurons. Therefore, disruption of STXBP1 function may lead to severe neurodevelopmental disorders^50^. However, in the patient reported in this study, no brain developmental or structural alterations were observed, thus suggesting the pathology is attributed to the direct function of STXBP1 on NT release, rather than to a gross brain development.

In order to provide a coherent interpretation of the clinical manifestation of R39dup and other mutations in the interface with syntaxin, we took advantage of *in vitro* studies including results from animal models^43^. In STXBP1 null mutant mice, NT release is completely blocked^51^. Recently, heterozygous Stxbp1+/− mice models were developed that recapitulate the seizure/spasm phenotype observed in humans^52^.

STXBP1 was implicated as an inhibitor and a stimulator of NT release across species^53–55^. Based on our analysis of the STXBP1-syntaxin 1A complex of the R39dup mutant relative to the WT, we suggest a schematic model of the clinical manifestation of the reported mutated protein (Figure 5). We show that STXBP1 R39dup exhibits a reduced affinity to syntaxin 1A. Thus, the binding affinity to syntaxin 1A in its closed, autoinhibited state is affected. Lowering the energy barrier (Table 2) for syntaxin 1A may switch syntaxin 1A to its open, extended state. Under such a conformational shift, the SNARE core domain of syntaxin 1A becomes accessible for interaction with the other SNARE proteins.

**Figure 4:**
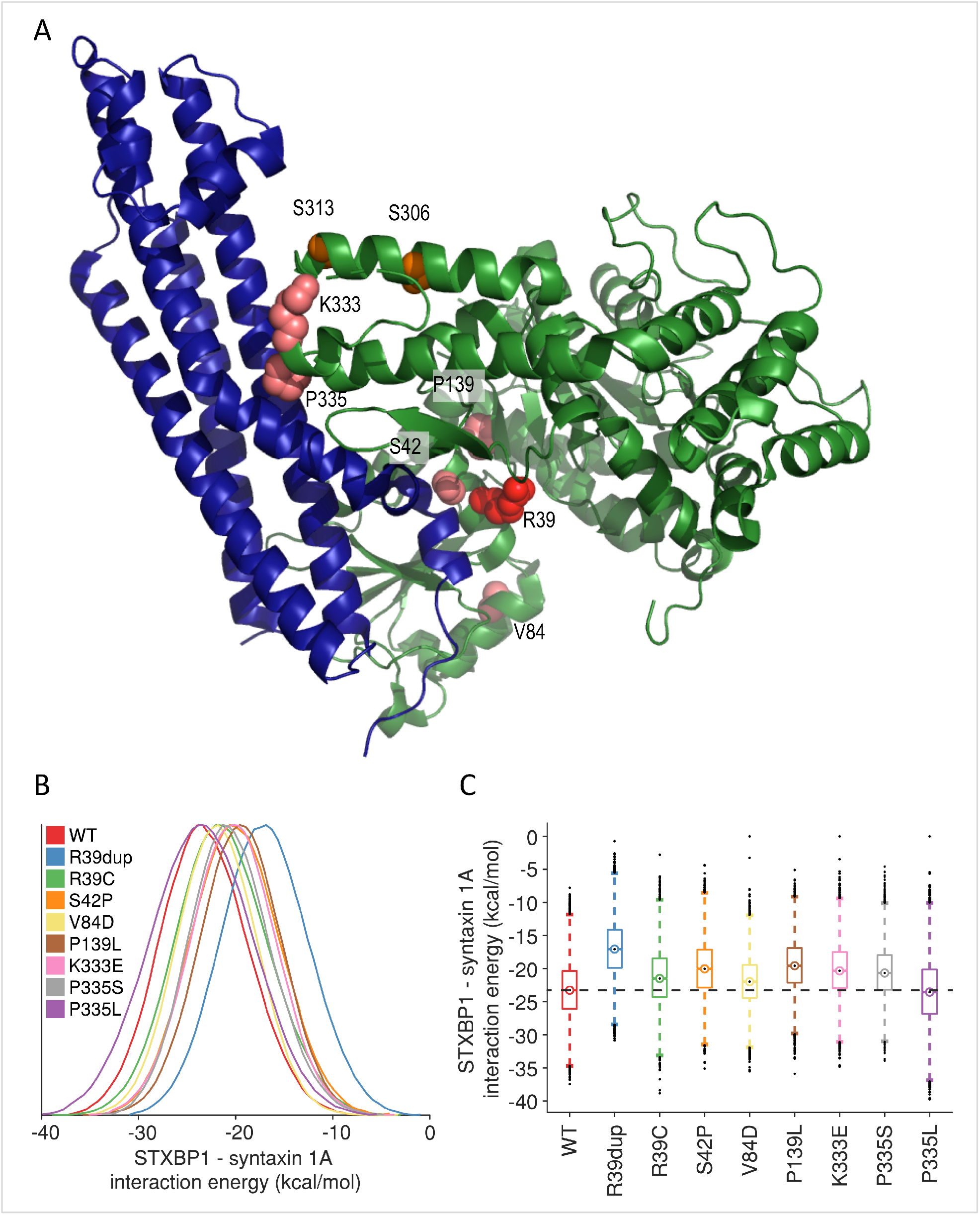
MD analysis of STXBP1 mutants, including the WT and the R39dup reported in this study. (A) Mutation site R39dup is indicated by a space fill colored red. The other 7 mutations are shown in a space fill, colored pink. All mutations are mapped to the STXBP1 - syntaxin 1A structure (PDB: 4JEH). (B) Histograms of FoldX interaction energies for STXBP1 - syntaxin 1A complexes (WT and mutants) calculated for the 10,000 simulation frames. (C) Box plots of the interaction energies for the trajectories frames. The center point is the median score, while 50% of the scores are within the box. The whiskers extend to cover >99% of the scores. The dashed line marks the interaction energies for the WT of STXBP1 - syntaxin 1A complex.

**Figure 5.**
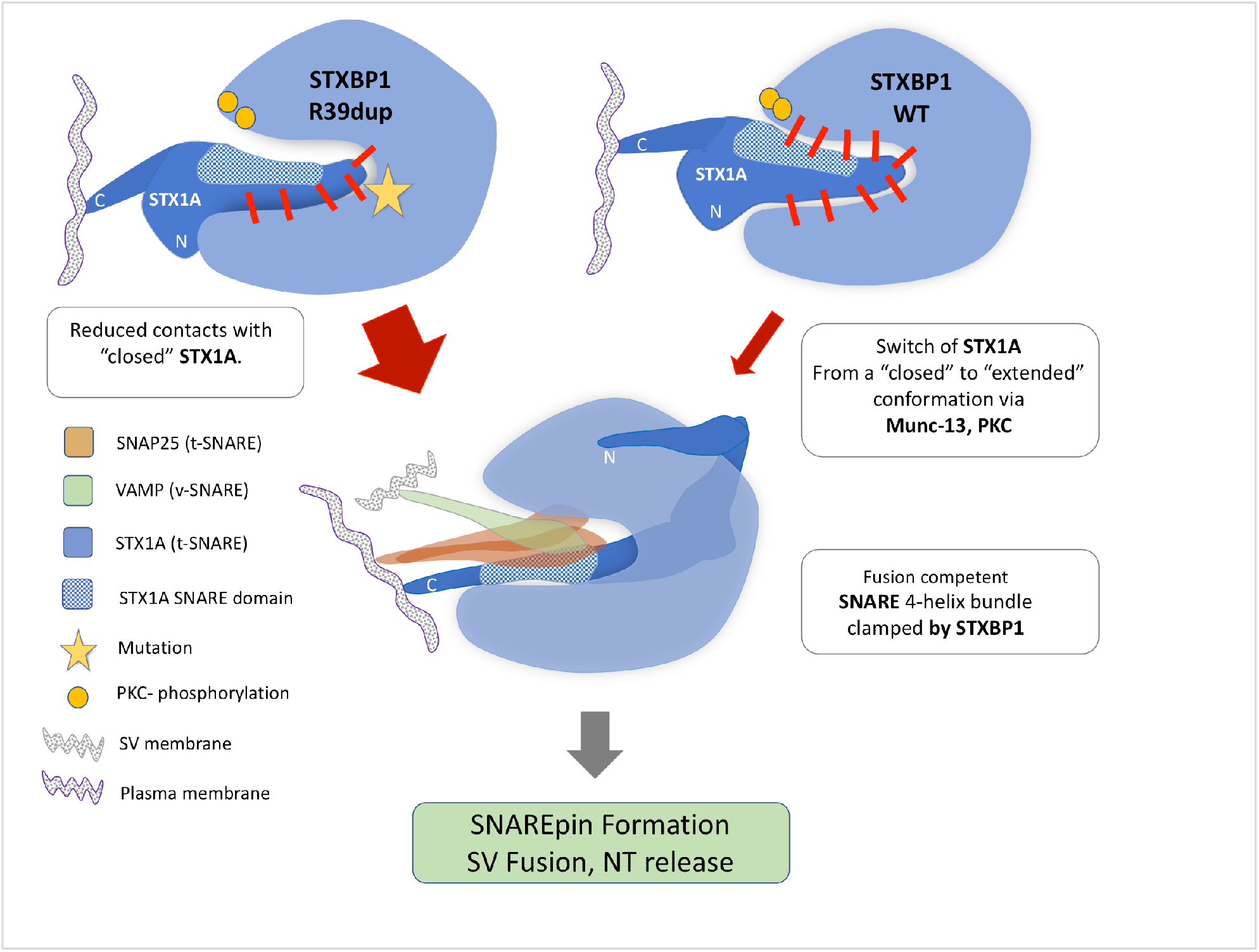
Model for STXBP1 pathology and clinical manifestation of R39dup mutant protein. A higher interaction energy of the R39dup mutant (yellow star, left) with respect to the WT protein complex of STXBP1-syntaxin 1A (right) results in a lower energy barrier for the syntaxin 1A transition from a closed, inactive state to an open, extended state. The strength of clamping of STXBP1 to syntaxin 1A is marked by the numbers of molecular clamps (short red lines). The switch to the extended state of syntaxin 1A exposes the SNARE domain of syntaxin 1A (Helix H3, dotted light blue pattern) that is involved in SNAREpin formation. The width of the red arrow symbolizes the higher probability of the SNARE domain of syntaxin 1A to engage in SNARE complex assembly. The sites of PKC phosphorylation are shown as yellow dots. Following phosphorylation, a process of formation of a tight 4-helix SNARE bundle is activated, leading to enhance membrane fusion and NT release. In the case of R39dup, the distance between the phosphorylations sites and syntaxin 1A was increased, presumably leading to a reduced dependency on phosphorylation for syntaxin 1A shifting to its extended, active state. The role of STXBP1 in supporting the SNAREpin formation is illustrated by showing the alternative binding mode of syntaxin 1A and STXBP1 and replacement of the closed syntaxin helices (Habc and H3) with the 4-helix SNARE bundle for SNAREpin formation, leading to a fusion of the SV membrane with the plasma membrane and NT release.

We further show that STXBP1 R39dup exhibits reduced contact with syntaxin 1A specifically at positions 280-338 of STXBP1 (Figure 2B). This segment overlaps with the two PKC-phosphorylation sites located at positions S306 and S313 (Figures 1C and 4A). These two sites are crucial for the direct interaction with syntaxin 1A^56^ and for NT release^57^. Specifically, it has been reported that PKC phosphorylation reduces the affinity for closed syntaxin 1A^43^, presumably with the assistance of Munc13-1 that causes a transition of syntaxin to the open, extended form. We propose that the reduced contact with syntaxin 1A at the PKC phosphorylation sites of the R39dup mutant reduces or completely relieves the PKC-dependency for switching syntaxin 1A from a closed to an open state.

The second phase in our model indicates the role of STXBP1 on NT release following the conversion of syntaxin 1A to the open state^58, 59^. Based on a detailed structure analysis and mutagenesis studies^60^, it was shown that the extended state of syntaxin 1A remains bound to STXBP1 via its N’-peptide (amino acids 4-24 of syntaxin 1A, remote from the R39dup site). This shift in the binding mode leads to a removal of the Habc helices of syntaxin from the central cavity of STXBP1. In such activated modes, STXBP1 acts as a platform, allowing VAMP that is a membranous protein of the SV membrane to activate the zippering of the SNARE 4-helix bundle^58, 59^. While the Habc helices of syntaxin 1A remain bound to the STXBP1 surface, the 4-helix bundle of the SNARE domain occupies the central cavity^53, 61^. STXBP1 presumably assists SNAREpin formation that creates the needed force for fusion of the SV membrane with the plasma membrane for executing NT release.

In sum, we demonstrate how an energetic shift in the interaction of STXBP1 R39dup with syntaxin 1A is converted to a change in the PPI dynamics that results in shifting the system towards NT release. Although we propose a molecular mechanism behind the actual impact of the mutation, we are still unable to correlate the mutation to the clinical phenotype. Specifically, we can only speculate why a high ΔΔG in this patient’s mutation (Table 2, Figure 4B and 4C) does not express as a severe epileptic encephalopathy in this patient. This merits further investigation and a data from cases of rare mutation in STXBP1, SNAREs and their direct interacting proteins.

Herein, we present a rigorous analysis for matching the clinical manifestation of patients with the direct genetic context. Medical geneticists may use such MD-based methodology for resolving inconclusive findings of variant inference and interpreting novel mutations. Resolving the mechanism of mutations is fundamentally based on structure, dynamics, and energetics of the protein within its natural context. Closing the gap between a report of a rare variant and the mechanism of action is critical for a correct diagnosis and accurate genetic counseling as well as for personalized therapy. The use of this method as a clinical tool is of particular importance, since current mutation effect prediction tools fail to predict the impact of protein interfaces and alterations in the network of protein interaction.

## Supplemental Data

Supplemental Data can be found online. Table S1 lists the results of mutation prediction effect tools. A structural shift is illustrated by anumation of the protein complex of STXBP-R39dup complexed with syntaxin1A.

## Acknowledgements

We gratefully acknowledge the patient families for their contribution in participation in this study. This study was supported by a fellowship to D.K (supported by Yad Hanadiv). We thank Prof Shy Arkin from the Hebrew University for providing us with high-performance computing resources to perform the MD analyses.

